# CUT&RUNTools 2.0: A pipeline for single-cell and bulk-level CUT&RUN and CUT&Tag data analysis

**DOI:** 10.1101/2021.01.26.428013

**Authors:** Fulong Yu, Vijay G. Sankaran, Guo-Cheng Yuan

## Abstract

Genome-wide profiling of transcription factor binding and chromatin states is a widely-used approach for mechanistic understanding of gene regulation. Recent technology development has enabled such profiling at single-cell resolution. However, an end-to-end computational pipeline for analyzing such data is still lacking. To fill this gap, we have developed a flexible pipeline for analysis and visualization of single-cell CUT&RUN and CUT&Tag data, which provides functions for sequence alignment, quality control, dimensionality reduction, cell clustering, data aggregation, and visualization. Furthermore, it is also seamlessly integrated with the functions in original CUT&RUNTools for population-level analyses. As such, this provides a valuable toolbox for the community.

## Introduction

Genome-wide analysis of transcription factor binding sites and chromatin states is essential for understanding cell-type specific transcriptional regulatory mechanisms. For over a decade, ChIP-seq has served as the main workhorse [1,2], but recently, a new generation of technologies has emerged with enhanced sensitivity and efficiency [3–8]. As a result, it has become possible to profile genome-wide occupancy analysis in a limited number of or even single cells. In the meantime, existing software packages do not have the capacity to analyze such data. New tools are needed to fill this important gap.

Among these technologies, CUT&RUN [3] and CUT&Tag [4] are the most popular. In previous work, we developed CUT&RUNTools for analyzing CUT&RUN data, providing an end-to-end CUT&RUN data analysis pipeline that includes sequence alignment and pre-processing, peak calling, cut matrix estimation, motif and footprinting analyses, and additional analyses [9]. Here we have further extended this software by implementing a flexible pipeline for single-cell data quality assessment, analysis and visualization, thus enabling users to rapidly utilize new technologies to systematically dissect the heterogeneity of the epigenomic landscape and gene regulatory networks among individual cells. In addition, we have also implemented a number of new features, including data normalization, peak calling, and downstream functional analysis that improve the performance for bulk data analysis. Together, this new tool is referred to as CUT&RUNTools 2.0, and is publicly available at https://github.com/fl-yu/CUT-RUNTools-2.0.

## Results

### Overview of CUT&RUNTools 2.0

CUT&RUNTools 2.0 provides a new module to facilitate the analysis and visualization of single-cell resolution data. The module implements a flexible, end-to-end pipeline that takes raw data as input, followed by a number of steps including data preprocessing and quality assessment, feature extraction, dimensionality reduction, cell clustering, data aggregation, and visualization. An overview of the single-cell pipeline is shown in Fig. 1. First, the input FASTQ files are processed by read trimming, mapping, and filtering. All the paired FASTQ files for individual cells are trimmed using a two-step strategy to improve the quality of the reads consistent with bulk data processing [9]. Then the trimmed reads are aligned to human/mouse reference genome using software Bowtie2 [10]. For each cell, only high mapping quality (MAPQ score > 30), uniquely aligned and properly mapped reads are retained for further analysis. CUT&RUNTools 2.0 uses GNU parallel technique [11] to improve computational efficiency in the main steps of data processing including reads trimming, mapping and filtering.

**Figure 1.**
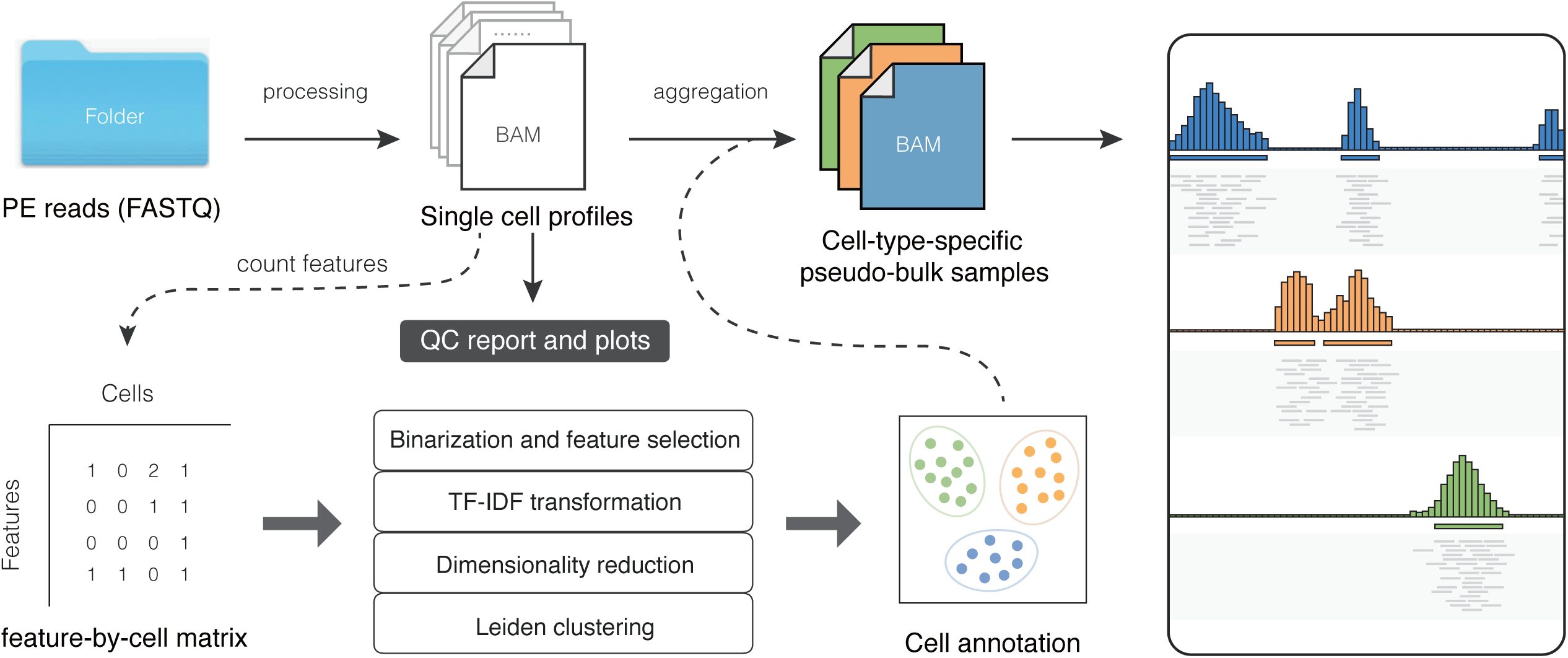
Overview of the workflow of CUT&RUNTools 2.0. The raw data of individual cells are trimmed and then aligned to the reference genome. Peak calling is performed for the aggregated data. A comprehensive QC report and several diagnostic plots are generated to help to evaluate experiment quality and filter low-quality cells. Three options of input features (*peak-by-cell, bin-by-cell* and *customFeature-by-cell*) are available for feature-by-cell matrix construction. The feature-by-cell matrix will be binarized and normalized, then the dimensionality reduction analysis is performed and the cell annotation will be generated from the clustering analysis. The cells from the same cluster are merged into a pseudo-bulk profile and the corresponding genome track files for both individual cells and the pooled signal are automatically generated for data visualization. The bulk-level analysis such as motif enrichment and Gene Ontology analysis provided by the CUT&RUNTools 2.0 could be used for functional annotation and interpretation of the resulting cell types.

CUT&RUNTools 2.0 reports a set of common quality control (QC) metrics as a summary report and diagnostic plots, which can be conveniently used for the data quality evaluation. In addition, single-cell level QC measures are saved and can be used to filter out low quality cells based on user-customized criteria. Furthermore, a signal-to-noise ratio metric is calculated based on the fraction of reads that fall into informative regions. To this end, all data of individual cells are pooled together into a single file followed by peak calling. The proportion of reads that fall into the detected peaks is defined as the signal-to-noise ratio in each cell.

Due to the sparsity of single-cell data, sequence reads falling into a set of pre-selected features are aggregated. CUT&RUNTools 2.0 provides three options for feature selection: peaks from cell aggregation, genome-wide bins, and user-defined functional elements. In each case, a feature-by-cell matrix is derived by counting the sequence reads that fall into a pre-identified feature across individual single cells in parallel. Furthermore, the count matrix is binarized to reduce noise associated with low-number counts. To efficiently reveal the latency variation across different cells, CUT&RUNTools 2.0 implements the term frequency-inverse document frequency (TF-IDF) transformation method to the binarized feature-by-cell matrix [12,13].

To reduce dimension, the resulting feature-by-cell matrix is processed by singular value decomposition, which generates a Latent Semantic Indexing (LSI) score matrix [12]. Then principal component analyses (PCA), t-distributed Stochastic Neighbor Embedding (t-SNE) [14], and/or Uniform Manifold Approximation and Projection (UMAP) [15] are used to further reduce dimensionality. Clustering analysis is achieved by applying the graph-based Leiden algorithm [16]. The cells from the same cluster are merged into a pseudo-bulk profile and the corresponding genome track files for both individual cells and the pooled signal are automatically generated per cell population. These pseudo-bulk samples are compared by analysis of distinct peaks, motif discovery, footprints (for transcription factors), or functional enrichment. The main processing steps in the data processing and feature-by-cell matrix construction can be performed in parallel to make full use of the available computational resources and reduce runtime. For each run, only a configuration file specifying the options and parameters needs to be provided for the analysis. Users can either run the entire workflow or select a specific step by customizing the configuration file.

### Analysis of a single-cell CUT&Tag dataset

To demonstrate its utility, we applied the CUT&RUNTools 2.0 pipeline to re-analyze a publicly available single-cell CUT&Tag dataset [4]. In this study, the investigators profiled genome-wide occupancy of H3K27me3, a repressive histone mark, in individual cells from two distinct cell lines: H1 (human embryonic stem cells) and K562 (a human erythroleukemia cell line).

A summary report regarding a set of QC metrics and the corresponding diagnostic plots for the experiment were produced (Fig. 2). Overall, a total of 1,373 cells were detected and approximately 0.14 million reads per cell were sequenced. For most cells, more than 99% of the reads were successfully mapped to the reference sequence indicating a high degree of purification. We also found that a vast majority of cells having a high proportion (median percentage, 99.5%) of nuclear reads (reads not aligned to mitochondrial DNA) in each single cell library. Less than 1% of duplicated reads were found for the majority of cells, suggesting the libraries of individual cells were sequenced near saturation. The fragment size was calculated as the length between the cut point of the Tn5 enzyme and the average size is 230.3 bp, which is expected for typical histone modification and longer than typical transcription factor binding profiles (∼120 bp) [4,9,17]. The fragment size distribution of all the reads from individual cells exhibits a clear nucleosomal binding pattern. These quality metrics were reported as a summary table (Fig. 2a) as well as a number of diagnostic plots (Fig. 2b and c). The high quality of the data is reflected by a number of factors including high alignment ratio, the ideal proportion of properly mapped reads, high-quality mapping reads and nuclear reads, and a high level of library complexity.

**Figure 2.**
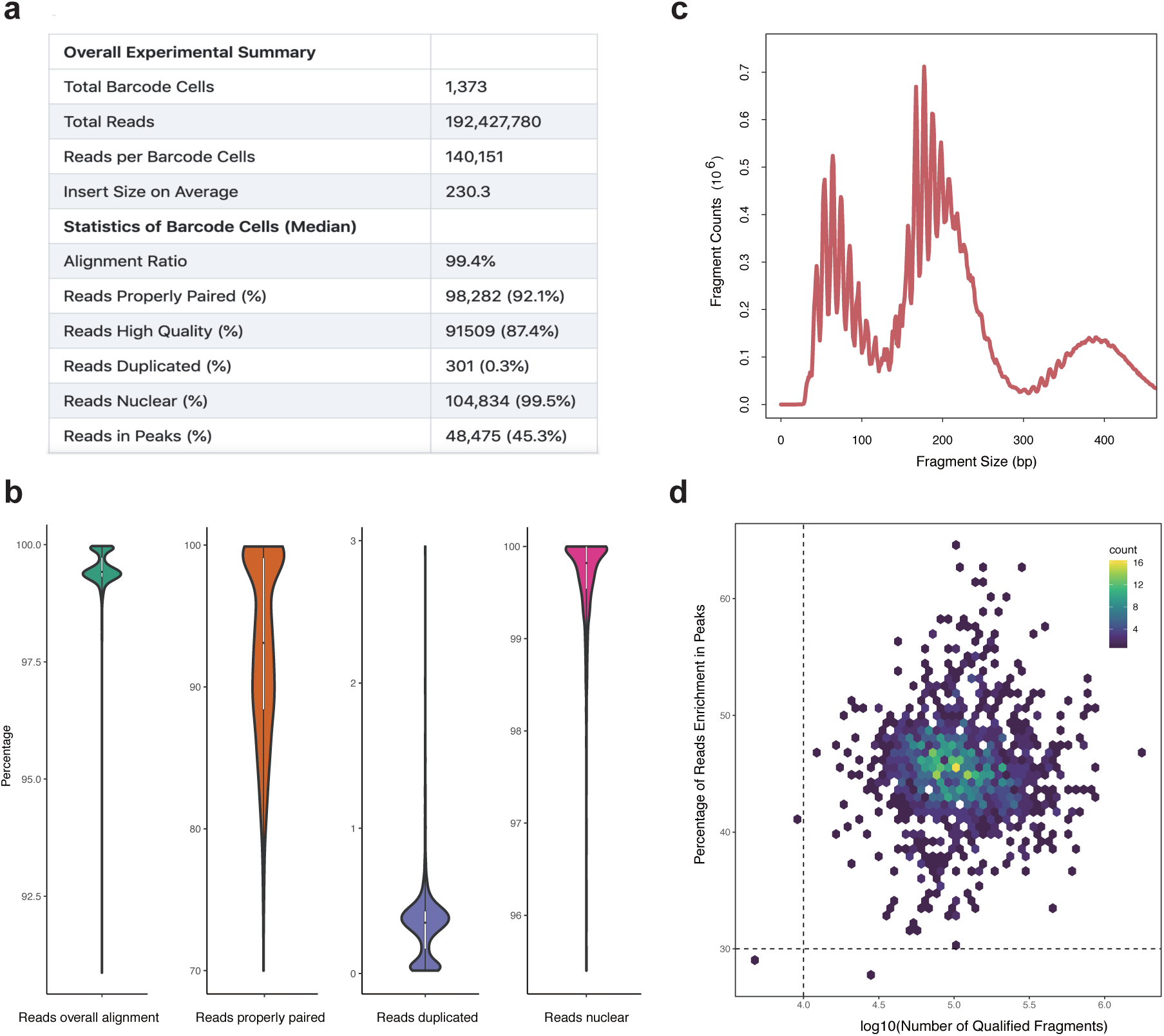
Quality evaluation of single-cell CUT&Tag data. (a) The overall statistics of the experiment are shown in the summary report. (b) Violin plots for percentage statistics of overall alignment, properly paired reads, duplication reads and nuclear reads. (c) Distribution of fragment size of the qualified fragments for all the cells. (d) Scatter plot of the fraction of unique fragments in peaks versus the total number of unique fragments for each cell. The default thresholds for the two parameters were indicated as dash lines.

Next, we aggregated sequence reads from individual cells into a pooled sample, and then applied MACS2 [18] to detect peaks. In order to preserve the structure of the data, we used a permissive cutoff of q-value < 0.01, which detects a total of 379,566 peaks. We assessed the signal-to-noise ratio in individual cells based on the fraction of reads that fall into the detected peaks. Overall, the signal-to-noise ratio ranges from 28% to 68%, with a median level of 45%. Of the 1,373 cells, three did not pass the QC criteria because they were associated with either a low signal-to-noise ratio (< 30%) or a small number of qualified fragments (< 10,000), therefore these three cells were excluded from further analysis (Fig. 2d).

For the remaining 1,370 cells, we created a binarized feature-by-cell matrix indicating the presence or absence of a peak of any individual cell. We also removed features that were either ubiquitous (detected in > 80% cells) or rare (detected in < 0.1% cells) therefore unlikely to be informative. After dimensionality reduction and clustering, two distinct cell populations were identified (Fig. 3a), which matched nearly perfectly to the true cell-type labels (Fig. 3b): nearly all the cells in cluster 1 were K562 cells, whereas all the cells in cluster 2 were H1 cells, indicating the biological information was preserved by our single-cell CUT&Tag analysis pipeline.

**Figure 3.**
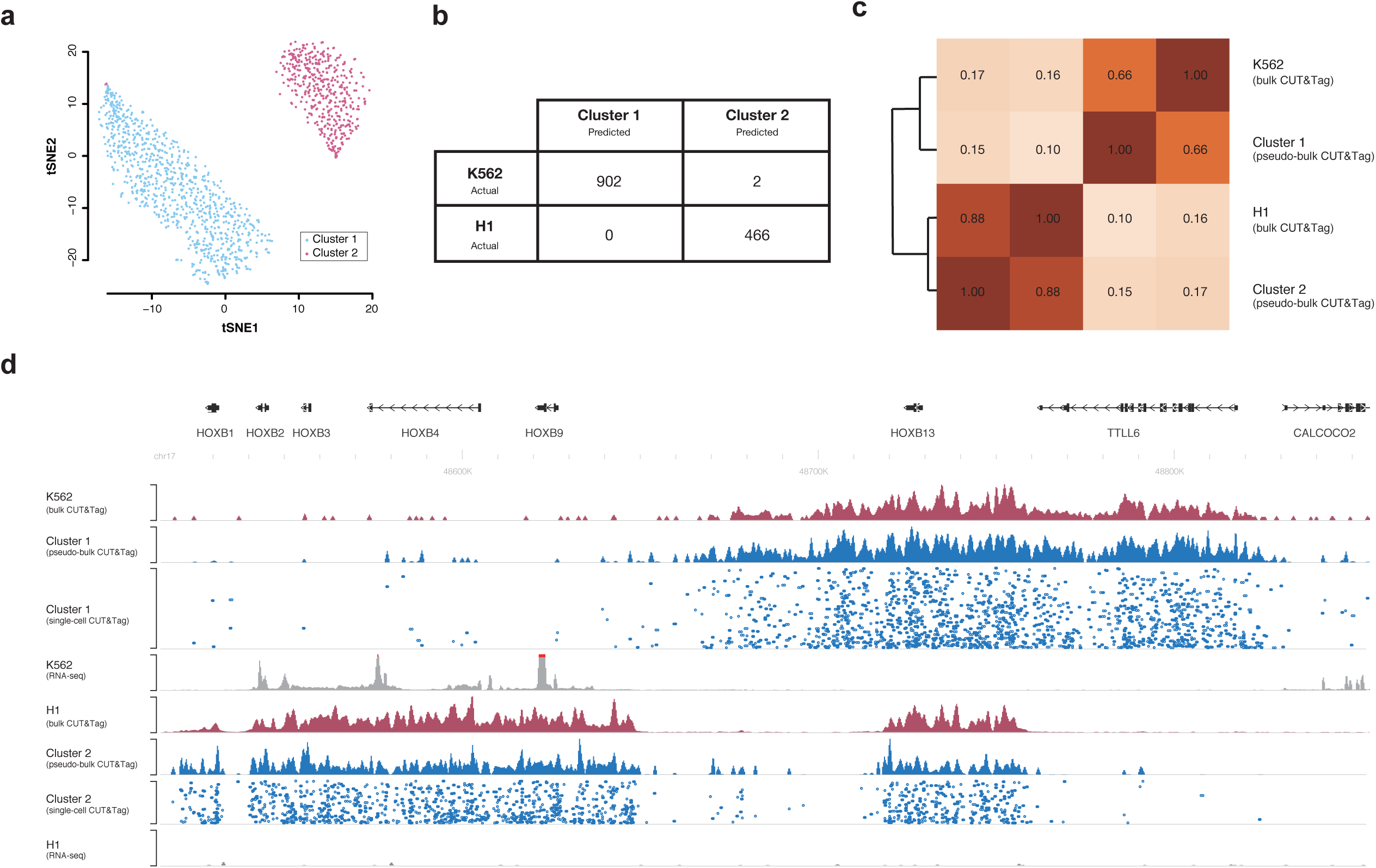
Analysis of single-cell CUT&Tag data identifies two cell clusters. (a) A plot of two-dimensional projection of the single-cell CUT&Tag data using the t-SNE method. (b) Confusion matrix of the ground truth cell labels and cell clusters predicted by CUT&RUNTools 2.0. (c) The genome-wide bins with 1 kb resolution were generated and the fragment within each bin was counted for the bulk and pseudo-bulk data of H1 and K562, respectively. The pair-wise Pearson correlation coefficients were calculated and shown in the heatmap. (d) The genome browser tracks for the HOXB gene locus. The signals of bulk, pseudo-bulk, and single-cell CUT&Tag data along with the RNA-seq data for H1 (top panel) and K562 (bottom panel) cells are shown. For the bottom panel, each row represents a single cell. The cells are ordered by total read counts.

To compare the genome-wide H3K27me3 profiles for different cell clusters, the reads obtained from all the cells in each cluster were aggregated to create a pseudo-bulk sample. We further downloaded and processed the cell-type matched bulk data and found the pseudo-bulk samples are highly correlated with the corresponding bulk data (Fig. 3c).

Together, these results suggest our single single-cell analysis is able to extract useful information and accurately reveal the cellular heterogeneity.

To aid visualization, we created genomic tracks files of not only the pooled signals, but also binding profiles at the single-cell resolution for different cell clusters (Fig. 3d). This visualization clearly shows the differences between the H1 and K562 cells. Of note, H3K27me3 occupies across the entire HOXB cluster in H1 cells, but only partially occupies a broad domain around the HOXB13 locus in K562 cells (Fig. 3d). By comparing with ENCODE RNAseq data, we found this change of H3K27me3 profiles is consistent with transcriptional activity differences between these two cell types, where HOXB1-9 genes are expressed in K562 cells but the entire HOXB cluster genes are repressed in H1 cells (Fig. 3d).

The pseudo-bulk data were used to further characterize and compare the H3K27me3 landscape between different cell subpopulations. We first identified 75,812 peaks in cluster 1 (corresponding to K562 cells) and 25,064 peaks in cluster 2 (corresponding to H1 cells) by using a stringent cutoff of q-value < 0.01 and fold change > 5 (Fig. 4a). We found only a small proportion of peaks (1,525) overlapping between these two clusters. More peaks were associated with non-coding regions comparing to coding regions in both cell clusters (Fig. 4b). Of note, a much larger proportion of peaks of cluster 2 (17%) were proximal to transcriptional start sites (TSSs) compared to cluster 1 (5%), suggesting that more embryonic associated genes may be more directly regulated by repressive H3K27me3 domain. We identified potential regulators closely related to the repression of cell-type-specific genes and cis-elements, such as the tumor suppressor Transcription Factor AP-2 Beta (TFAP2B) and Early B cell factor 1 (EBF1) in cell cluster 1 [19,20] (Fig. 4c) and the development associated TF Early growth response protein 2 (EGR2) in cell cluster 2 [21,22] (Fig. 4d). Gene Ontology analysis showed that many different cell and system development associated functions including embryo development, system development, cell differentiation, and multi-cellular organism development were markedly enriched in cluster 2, which also supports that the establishment and removal of H3K27me3 at specific genes in the embryonic stem cells is critically important for normal development.

**Figure 4.**
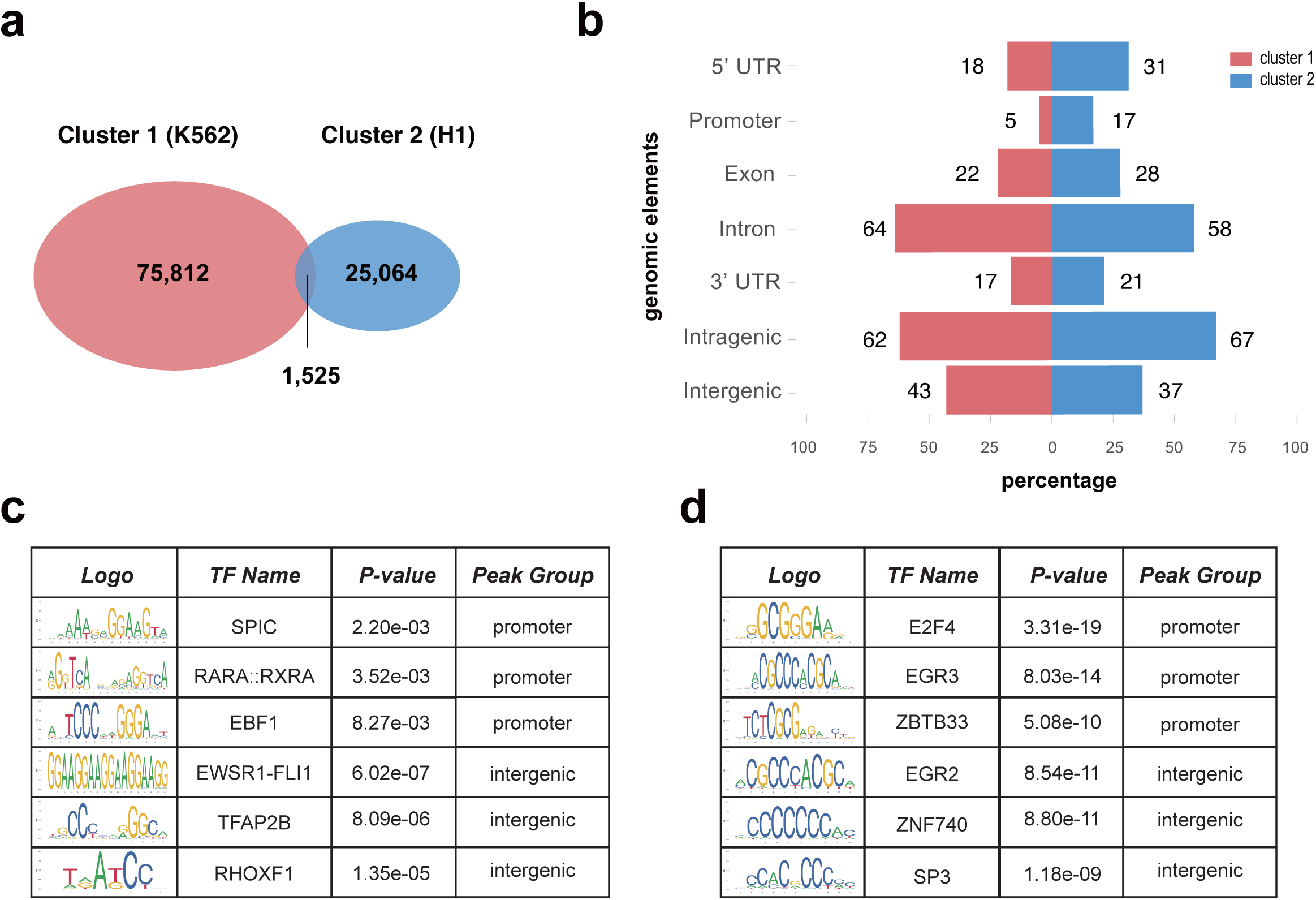
Functional analysis of the pseudo-bulk data. (a) Venn diagram illustrates the number of H3K27me3 modification peaks detected in the pseudo-bulk data corresponding to the two cell populations. (b) The percentage distribution of top 5000 peaks overlapping with annotated genomic elements. (c-d) The top enriched TFs associated with different genomic elements of cell cluster1 (c) and cell cluster 2 (d).

## Discussion

We have extended CUT&RUNTools in response to the recent development of single-cell technologies and demonstrated that this new single-cell module is useful for extracting biological insights. While we have only focused on single-cell CUT&Tag here, our pipeline can be adapted to various emerging promising technologies that could profile transcription factor and chromatin regulators at the single-cell resolution [5–8,23]. As more data with a large sample size become available, we plan to incorporate further adaptations to this approach in the future.

In addition, CUT&RUNTools 2.0 also contains a number of updates in bulk-level analyses, such as spike-in sequence alignment and data normalization (see the website for details). The main strength of CUT&RUNTRools 2.0 is that it seamlessly integrates single-cell and bulk-level analyses in one package, providing the convenience to study a number of datasets in a standardized manner. As single-cell multi-modal data become increasingly available, CUT&RUNTools 2.0 provides as a convenient toolkit facilitating integration which in turn will provide a better understanding of epigenomic heterogeneity and regulatory logic in both healthy and diseased tissues.

## Methods

### Adaptor trimming and reads mapping

For each cell, the adaptor sequence and primer oligo sequence from the 3’ ends of reads is trimmed off using a two-step trimming strategy consistent with bulk data processing. All the reads are aligned to the corresponding reference genome hg38 using Bowtie2 software [10] with the default parameter. The SAMtools [24] is used to sort and index the resulting BAM files.

### Quality control metrics

CUT&RUNTools 2.0 provides a set of quality assessment metrics for the overall experiment as well as each barcode cells. The metrics include overall alignment ratio, the number of total reads, properly paired reads, duplicated reads, high-quality reads (MAPQ score >30), and nuclear reads, fragment size, signal-to-noise ratio (the fraction of reads in peaks). The Picard (https://broadinstitute.github.io/picard) *MarkDuplicates* function is used to locate and tag duplicate reads. The unmapped, low quality (MAPQ score <30), unproperly paired and duplicated fragments are discarded, and the remaining data are defined as the qualified fragments. Then, the filtered and sorted BAM file for each cell are generated for further analysis. The fraction of reads in peaks are calculated using BEDTools *genomecov* function [25].

For each run, a summary report is generated by using a custom script, which contains a summarization of the mapping statistics of the overall experiment. Additionally, several diagnostic plots are also produced for intuitive illustration of the quality control metrics across all the cells. We provide two parameters, *num_reads_threshold* (number of unique mapped reads, 10,000 as default) and *percentage_rip* (fraction of reads in peaks, 30% as default) as filters to remove low quality cells. Finally, two files, *statistics_QCpassed*.*txt* and *statistics_QCfailed*.*txt*, are generated to record the identity of the filtered cells along with their associated statistics.

### Parallel processing

Comparing to bulk data, a single-cell CUT&RUN / CUT&Tag experiment usually contains a large number of cells, but the reads in each cell are less abundant. The data processing time roughly scales linearly with the cell number, therefore it may take a long time if the number of cells is large. To overcome this challenge, we have adopted the GNU parallel technique [11] in the main steps of data processing, which results in dramatic reduction of runtime.

### Peak calling

For the pseudo-bulk data aggregated from cells from the entire datasets or a specific cell cluster, CUT&RUNTools 2.0 enables peak calling by using different methods. By default, we use the MACS 2 narrow peak mode for peak calling [18], which has good performance for TFs [26]. In addition, we also implement two alternative strategies: the MACS 2 broad peak mode, and the SEACR algorithm [27]. The users can easily make selection by modifying the parameters of *peak_caller* and *peak_thresholds*.

### Construction of feature-by-cell matrix

CUT&RUNTools 2.0 provides three options to build the feature-by-cell matrix, which can be customized by setting *matrix_type* as *peak-by-cell, bin-by-cell, or customFeature-by-cell. peak-by-cell* refers to using peaks detected from the pooled sample as the feature set, which is selected in the present study. *bin-by-cell* refers to segmenting the genome into equal-size bins (5 kb by default). Using the *customfeature-by-cell* option, users can also upload their own features of interest, such as a list of enhancer regions (input needs to be in the standard BED format).

Once the feature file is designated, CUT&RUNTools 2.0 automatically excludes features overlapping with ENCODE blacklist regions [28] or uninterested chromosomes. For CUT&RUN experiments, an additional filtering step is carried out by removing the regions overlapping with TA repeats regions because these regions usually occur as containment abnormally enriched reads [17]. The users can simply set the *experiment_type* parameter as CUT&RUN to turn on this feature. With the preparation of feature files, CUT&RUNTools calculates the read coverage profiles for all the qualified cells, and the resulting feature-by-cell count matrix is generated and saved.

### Count matrix processing

Owing to the sparsity of count matrix of single-cell epigenomic data, CUT&RUNTools 2.0 converts the feature-by-cell matrix into the sparse Matrix format to allows more efficiency of memory usage and computation by using the ‘Matrix’ package in R [29]. The sparse matrix is binarized and an additional filter is used to remove features that are present only in few cells (the maximum of 0.1% of the cells) or in the vast majority of cells (80% of the cells) to efficiently capture the informative signals.

### Dimensionality reduction, cell clustering and visualization

Dimensionality reduction is performed using the Latent Semantic Indexing (LSI) method, a technique commonly used for document indexing process in natural language processing [30]. The binarized sparse matrix is first converted into a frequency-inverse document frequency (TF-IDF) matrix by weighting the matrix against the total number of terms (i.e. features) for each document (i.e. cell) with the following formula:

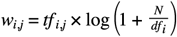

where *w*_*i,j*_ is the weight for the feature *i* in cell *j, tf*_*i,j*_ indicates the term frequency, the number of feature *i* in cell *j, df*_*i*_ is the document frequency of term *i*, cell number of cells where the feature *i* appears, *N* is the total number of cells in the experiment.

The singular value decomposition (SVD) is applied on the TF-IDF matrix *X* to generate an LSI score matrix with a lower k dimensional space as follows:

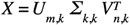

This is the decomposition of *X* where *U* and *V* are orthogonal matrices and ∑ is a diagonal matrix. *m* represents the number of rows and *n* represents the number of columns for *X. U* = [*μ*_1_,…, *μ*_*k*_] and *μ*_*i*_ with length *m*, which is called the left singular vector. *V* = [*v*_1_,…,*v*_*k*_] and *v*_*i*_ with length *n*, which is called the right singular vector. *∑* _*k*_,_*k*_ = *diag* (*σ*_1_,…, *σ*_*k*_) and *σ*_1_≥ *σ*_2_≥ *σ*_*k*_ are singular values of *X*.

Based on the resulting LSI matrix, the graph-based Leiden algorithm is used as the unsupervised clustering method which is implemented with the *leidenalg* and *igraph* library [16,31]. Using the top 30 principal components (different numbers can be tuned by changing parameter *cluster_pc*), we build a shared neighbor network (SNN) graph by considering each cell as a node and further finding its k-nearest neighbors according to the Euclidian distance. Another parameter of *cluster_resolution* is also provided, which is important to control the number of resulting clusters, where the larger of this parameter usually lead to the larger number of resulting cell clusters. The predicted cell type labels are then generated. CUT&RUNTools 2.0 visualizes the results by employing three commonly used methods for dimensionality reduction including PCA, UMAP and t-SNE. The corresponding plots of the first two dimensions are generated and saved, respectively.

### Generation of genome browser tracks

The tracks of pseudo-bulk data are normalized by the counts per million method and generated with bigwig format using the program *bamCoverage* in the deepTools package [32] with 50 bp resolution. The peak files called from pseudo-bulk data are also generated with BED format. For each single cell, the genome-wide read coverage is calculated using BEDTools [25] in parallel with available computational cores. The regions covered by any reads are extracted and a read coverage file is created with BED format. For each cell cluster, the read coverage files of individual cells are combined into a single track with qBED format for convenient visualization [33], with the cells sorted by the number of covered regions in each cell. All the resulting track files could be directly uploaded and visualized using browser apps such as the WashU Epigenome Browser [34].

### Downstream analysis function specific to cell populations

Analysis of peaks called from pseudo-bulk data of cell clusters is important to uncover the cell-specific gene regulatory elements. The script *peakOverlap* is used to summarize the peak overlap between different cell clusters, the number of common and specific peaks will be identified and visualized as a Venn diagram. The script *eleAnno* is used to annotate the distribution of peaks over different types of genomic features including 5’ UTR, promoter, exon, intron, 3’ UTR, intragenic and intergenic regions. *haystack_motifs* is used for the motif enrichment analysis of peaks to identify cell-type-specific cis-regulatory elements and associated transcription factors [35]. *PeakFun* is used for Gene Ontology (GO) analysis of the top 1,000 interested peaks with the search of genes associated with GO ‘biological process’ categories [36] (see the manual on the website for details).

## Declarations

## Ethics approval and consent to participate

Not applicable.

## Consent for publication

Not applicable.

## Availability of data and materials

We applied CUT&RUNTools 2.0 to a single-cell CUT&Tag dataset available on SRA database (SRP190015 and SRP175327). For the purpose of comparison between bulk and single-cell data, raw FASTQ data of H3K27me3 bulk CUT&Tag for H1 (SRX5193360) and K562 (SRX5193370) were downloaded and processed using CUT&RUNTools 2.0 bulk-data analysis pipeline. The RNA-seq data for H1 (ENCBS734AAA) and K562 (ENCBS864OKZ) were downloaded from the ENCODE website. The genomic annotation data were downloaded from UCSC table browser. The CUT&RUNTools 2.0 software is freely available under the MIT license.

## Competing interests

The authors declare that they have no competing interests.

## Funding

This work was supported by an NIH grant R01HG009663 to G-C.Y and grants from New York Stem Cell Foundation and NIH (R01DK103794) to V.S.

## Authors’ contributions

F.Y. and G-C.Y. conceived and designed the study. F.Y. designed and implemented the single-cell processing module, F.Y. designed and implemented the new functions for bulk data analysis. G.-C.Y. supervised the overall study. F.Y. and G.-C.Y. wrote the paper with input from V.S. All authors read and approved the final manuscript.

## Acknowledgements

We thank Dr. Qian Zhu for providing helpful advice and feedback for this paper.

## Notes

### Competing Interest Statement

The authors have declared no competing interest.

